# Impact of cancer mutational signatures on transcription factor motifs in the human genome

**DOI:** 10.1101/422386

**Authors:** Calvin Wing Yiu Chan, Zuguang Gu, Matthias Bieg, Roland Eils, Carl Herrmann

## Abstract

**Background:** Somatic mutations in cancer genomes occur through a variety of molecular mechanisms, which contribute to different mutational patterns. To summarize these, mutational signatures have been defined using a large number of cancer genomes, and related to distinct mutagenic processes. Each cancer genome can be compared to this reference dataset and its exposure to one or the other signature be determined. Given the very different mutational patterns of these signatures, we anticipate that they will have distinct impact on genomic elements, in particular motifs for transcription factor binding sites (TFBS).

**Results:** In this work, we build the link between mutational signatures and TFBS motif alterations. We investigated and computed the theoretical impact of mutational signatures on 512 TFBS motifs, hence translating the trinucleotide mutation frequencies of the signatures into alteration frequencies of specific TFBS motifs, leading either to creation of disruption of these motifs. We further build a theoretical prediction of the alteration patterns for different cancer types based on the exposure of these cancer types to the mutation signatures. For certain motifs, a high correlation is observed between the TFBS motif creation and disruption events related to the information content of the motif.

**Conclusion:** Our results show that the mutational signatures have different impact on the binding motifs of transcription factors and that for certain high complexity motifs there is a strong correlation between creation and disruption, related to the information content of the motif. This study represents a background estimation of the alterations due purely to mutational signatures in the absence of additional contributions, e.g. from evolutionary processes.

## INTRODUCTION

With the availability of thousands of fully sequenced cancer genomes, genome-wide patterns of somatic mutations can be analyzed to search for potential driver mutations. Such an effort has been exemplified by the recent PanCancer Analysis of Whole Genomes initiative by the International Cancer Genome Consortium (ICGC) consortium. However, beside coding driver mutations that have been described earlier, non-coding mutations have been under increased scrutiny, in search for additional non-coding drivers, given the extensive number of these mutations in non-coding genomic regions [1, 2]. Several modes of actions can be identified for these mutations, the most likely ones being mutations affecting regulatory elements such as transcription factor binding sites, altering in one way or the other (creation or disruption) the binding motif. A spectacular example was identified in several cancer entities involving the promoter of the TERT gene, in which two recurrent mutations have been shown to create new binding sites for ETS-family transcription factors, leading to a strong over-expression of the TERT oncogene [3, 4]. Another example was found in T-ALL, in which a mutation creating a new binding site for MYB leads to the appearance of a super-enhancer driving over-expression of the TAL1 oncogene [5]. Besides these few individual obvious non-coding drivers, cancer genomes are loaded which thousands of mutations which are termed passenger, as they cannot be individually related to molecular phenotypes, as in the previous cases. However, several studies have shown that these putative passengers contribute to an overall mutational load in the cancer genome, and can, collectively, have an impact [2, 6]. Hence, it is of importance to understand the overall patterns of non-coding mutations, besides the few driving examples.

Patterns of somatic mutations have been analyzed by defining so-called *mutational signatures*, based on a dimensional reduction approach focusing on the patterns of trinucleotide alterations. The 96 possible types of trinucleotide mutations were summarized into a reduced number of signatures, which each describe a different mutational bias [7]. Some of these mutational signatures can be related to specific mutational processes such as APOBEC mutations, nucleotide mismatch repair or various carcinogens. Once these overall signatures are available, the exposure of cancer types or of individual cancer genomes can be determined. Hence, for example, there is a clear association between the signature related to ultra-violet light and melanoma [8].

In study, we assess the impact of mutational signatures on motifs of transcription factor binding sites. How does a particular mutational signature impact the large collection of transcription factor binding site motifs? We want to establish a catalogue, for each signature, of binding motif creation and disruption frequencies, which would correspond to an expected background effect of the mutational patterns, in the absence of any additional effect such as selective processes. Our goal is thus to translate the mutational signatures based on trinucleotide alterations into signatures on motif alteration. The result provides a theoretical framework as a baseline model for transcription binding site alteration analysis in cancer genomes.

## METHODS

The following regulatory impact analysis is based on the 30 trinucleotide mutational signatures described by Alexandrov et al. [7]. These mutational signatures were downloaded from the COSMIC database (https://cancer.sanger.ac.uk/cosmic/signatures). Each of the 30 mutational signature contains the normalized mutational probability across the 96 types of point mutation in a trinucleotide context. In the remaining of this paper the mutational signature is represented by the 30 x 96 mutational signature matrix *S*_*M*_ and mutational signature vector 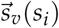 as follow:

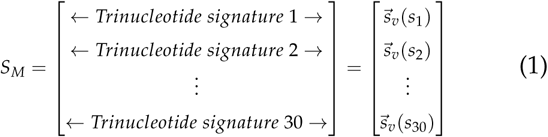

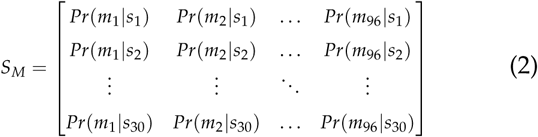

where *m*_*i*_ corresponds to the 96 possible trinucleotide mutations (eg. A[C>A]A, A[C>G]A,…etc.). This mutational signature matrix is denoted as the *trinucleotide mutational signature* in the remaining part of this article.

### Transcription Factor Motif Alteration Signature

The procedure of computing the motif binding alteration probability is illustrated in Figure 1a.

**Figure 1.**
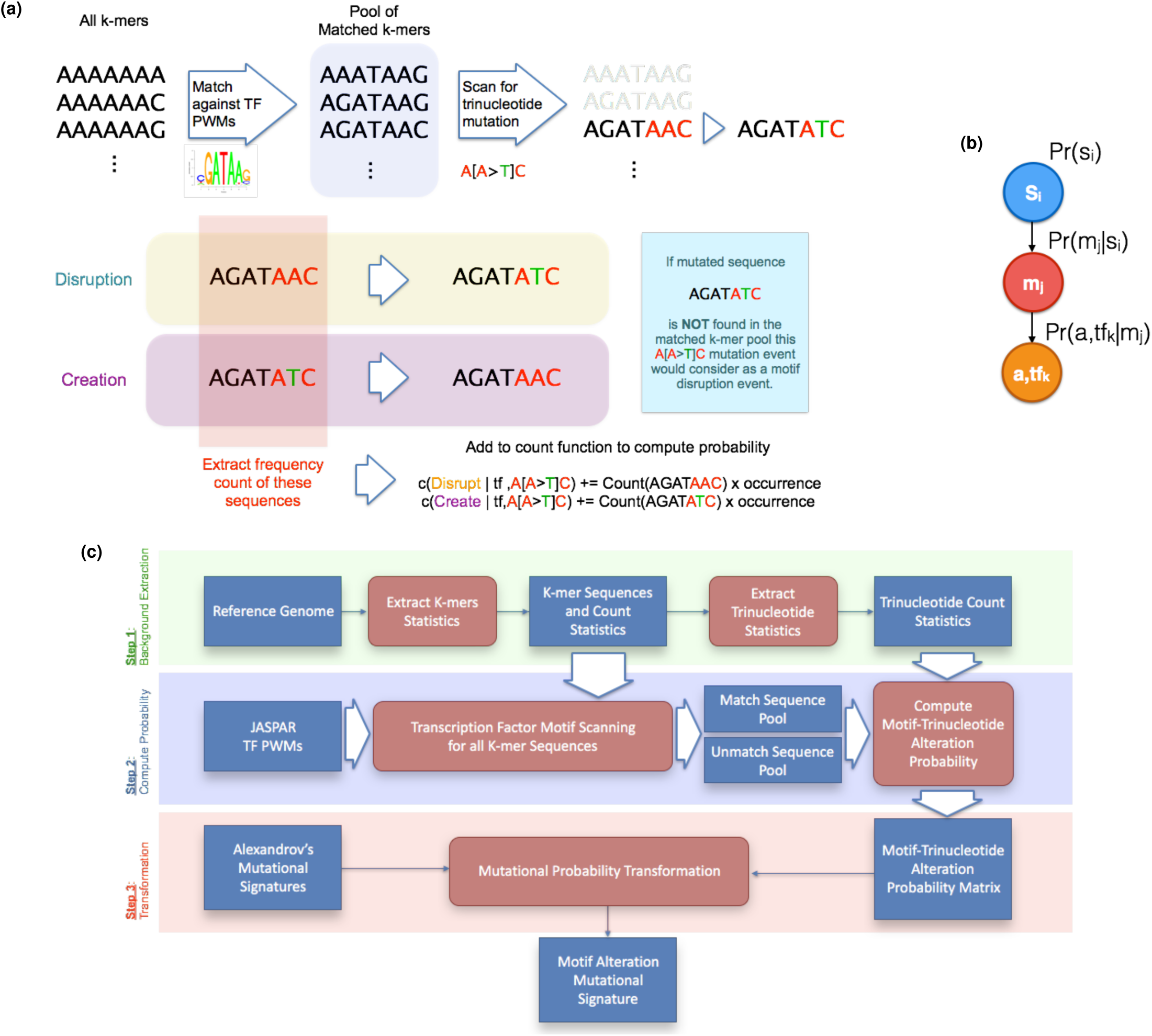
Computing motif binding alteration probability: (a) Frequency counting procedure; (b) Bayesian tree representation of joint probability; (c) Transcription factor binding motif alteration signature analysis workflow.

We define *Pr*(*a | t f*_*i*_, *m*_*j*_) as the probability for a motif *t f*_*i*_ to undergo an alteration *a* (i.e. creation or disruption) given a trinucleotide mutation *m*_*j*_. A disruption event is defined as a mutation that turns a binding site into a non-binding site, and a creation event corresponds to the reverse effect. The probability of a transcription factor motif alteration event is computed by assessing the p-value of a given sequence before and after the mutation. The 512 JASPAR 2016 vertebrate position weight matrix (PWM) of length 6 to 19 are used to compare the impact in binding affinity due to a point mutation [9]. The p-value of a binding site is evaluated using the matrix-scan tool of the Regulatory Sequence Analysis Tools toolbox to compute the p-value [10]. In this study, a pvalue of p=0.001 is set as the binding threshold. For a given transcription factor binding PWM of width *k*, all possible k-mer sequences are scanned to compute the mutational statistics. For a PWM of *k* = 19, it requires scanning a total of 4^19^ = 274, 877, 906, 944 for all the 19mer sequences. The search space of binding sequences can be reduced in half by combining all the reverse complementary sequences.

For each PWM *t f*_*i*_ of width *k*, we enumerate all possible k-mers (considering a k-mer and its reverse complement as the same motif) and, using matrix-scan with the parameters described above, we separate the set of k-mers into binding and non-binding k-mers. Then, the alteration probability is computed by mutating each of the trinucleotide in the matching k-mer according to the mutation type *m*_*i*_ and search for the corresponding mutated sequence. A disruption event is identified if the mutated sequence is not found in the list of binding k-mers. Conversely, a creation event is identified if a non-binding k-mer is turned into a binding k-mer. The current matched binding sequence is considered as the reference sequence for a motif disruption event and alternative sequence in the motif creation event and vice versa.

In order to count these events in the human genome, all possible k-mers are extracted from the human genome (version hg19) and their occurrences are counted for *k* = 6 to *k* = 19. The count of each of the reference sequence in the human genome is recorded according to the type of trinucleotide mutation and the type of alteration event. The probability *Pr*(*a| t f*_*i*_, *m*_*j*_) can be obtained by normalizing the counts by the total number of the reference trinucleotide *c*_*hg*19_(*m*_*j,re* *f*_) for the reference trinucleotide *m*_*j,re* *f*_ of a given mutation type *m*_*j*_ of a *t f*_*i*_ PWM width of length *w*(*t f*_*i*_). The count normalization factor for alteration probability computation is illustrated in Figure 2. For each trinucleotide in the genome, it is compared *w*(*t f*_*i*_) *-*2 times. Therefore, the total count detected should be divided by the number of trinucleotides in the genome multiplied by *w*(*t f*_*i*_) *-* 2 to obtain the alteration probability

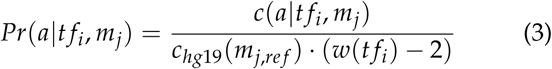

**Figure 2.**
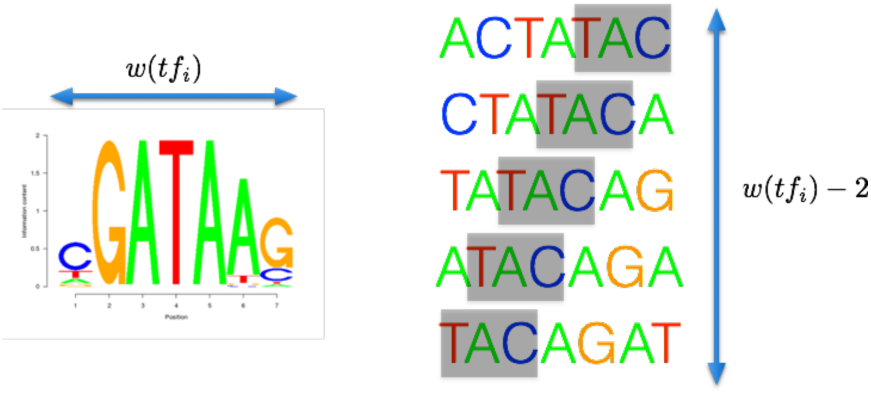
Illustration of alteration counts normalization factor. Each trinucleotide in the reference genome is scanned *w*(*t f*_*i*_) − 2 times with the TF motif of length *w*(*t f*_*i*_).

It is important to note that the conditional probability is computed with *t f*_*i*_ given to ensure that the number of motifs in the database does not influence in the probability. Importantly, we also need to determine the binding affinity of k-mers which do not occur in the human genome for the motif creation event, as a mutation could turn a k-mer into one which does not occur in the reference genome. However, for the disruption probability computation, the k-mers search space for genomic k-mers count can be drastically reduced by considering only k-mers occurring in the human genome, which dramatically reduces the search space for *k >* 13. The motif alteration probability of a given transcription factor PWM and trinucleotide is stored in the motif alteration probability matrix,

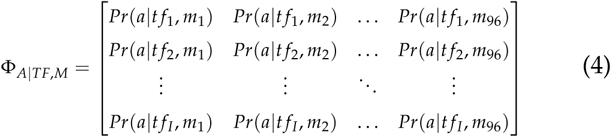

From this, we compute the alteration probability for a mutational signature *s*_*i*_ using Bayesian inference:

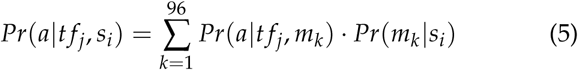

or, in matrix notation:

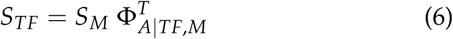

where,

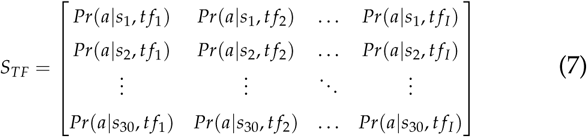

The workflow of the transcription factor motif alteration signature is shown in Figure 1c and the complete algorithm for computing the alteration probability is given in the Appendix.

### Analysis on Transcription Factor Motif Alteration Probability

The motif alteration probability matrix Φ_*TF|M*_ encapsulates changes in binding affinity of a given PWM under the perturbation of a single nucleotide point mutation. In order to investigate similarities in the alteration probabilities of different motifs, a hierarchical clustering of motif creation and motif disruption is performed. A Partitioning Around Medoids (PAM) clustering approach is applied to partition the 512 transcription factors using the silhouette coefficient to determine the optimal number of groups. The clustering is then compared to the TF family annotation downloaded from the JASPAR database. To further investigate the alteration probability of relevant TFs in cancer, all transcription factors present in the COSMIC cancer gene census are extracted and evaluated. The global alteration offset of these COSMIC cancer TFs are computed by subtracting the disruption probability from the creation probability.

The hierarchical clustering provides a tree structure of paired similarity between the set of input labels (TF motifs), and the PAM clustering results in a strict partition of the motifs. These methods are less than ideal for visualizing the clustering and distance at the same time particularly when each of the TF motifs is described by a high dimensional vector (192 for trinucleotide alteration probability and 60 for motif alteration signatures). To evaluate the similarity of motif alteration probability of multiple transcription factors, a self-organizing map (SOM) analysis is performed using the alteration probability matrix Φ_*TF*_ *|* _*M*_ to validate the result and to gain insight into the alteration similarity among the TF motifs. For this, a 22 x 22 grid was used and resulted in a stack of 96 variable maps corresponding to the trinucleotide mutation type *m*_*k*_. The map dimension of the SOM is selected to maximally retain the resolution of the transcription factor space based on the following criterion:

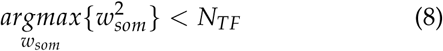

where *w*_*som*_ is the width of the SOM, and *N*_*TF*_ is the total number of TFs. There is a total of 512 TFs in the JASPAR vertebrate 2016 database, therefore the optimal SOM dimension is *w*_*som*_ = 22. This allows the TFs to be distributed evenly on the map in the worst case scenario when they are equally dissimilar with respect to each other.

### Comparison with the TCGA Dataset

We compared our motif alteration prediction based on the mutational signatures with real datasets of SNVs in cancer. For this, we used a previously published TCGA dataset with 505 samples covering 14 cancer entities to validate the results [11].

In order to predict motif alterations in the TCGA dataset, we developed a motif alteration pipeline. The pipeline is based on the matrix-scan-quick tool from the RSAT toolbox [10]. The neighbourhood reference sequence of each SNV is extracted to build a second order Markov model for PWM matching, both the reference and alternative sequence are matched against the given PWM. For alteration detection, it is required to fulfill both of the following conditions: (1) the mutation results in a change from binding site to non-binding site (or vice-versa for creation) where a threshold of *p ≤* 1*e -* 4 is required to consider a sequence as a TF binding site; (2) a 10 fold p-value change before and after the mutation.

The TCGA SNV calls are first matched against the same set of JASPAR PWMs to identify possible transcription factor binding alteration sites. To produce comparable results across all cancer entities, the detection probability of each of the transcription factor alteration is computed by normalizing the detection counts with respect to the total number of SNV detected in the corresponding cancer entity.

To match our predicted frequencies of alteration based on the mutational signature to the cancer dataset, we need, for each of the cancer entities, to combine the influence of those mutational signatures that contribute to the particular cancer entity. Here, we used, for each cancer sample, the exposure matrices described in [12], based on 482 (out of the 505 samples in [11]) overlapping TCGA samples. To infer the transcription factor binding site alteration probability for one cancer entity, the transcription factor motif alteration matrix *S*_*TF*_ is multiplied with the normalized exposure matrix *E*_*PID*_ to produce the per patient exposure prediction matrix Ψ_*PID*_,

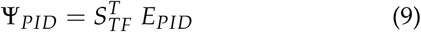

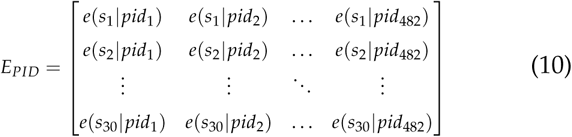

where *e*(*s*_*l*_ *| pid*_*m*_) is the normalized exposure (or exposure probability) of signature *l* and patient *m*. Combining the above we have,

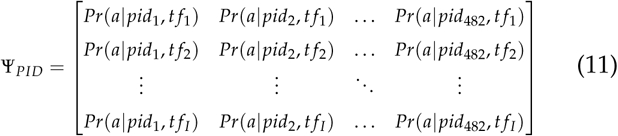

After obtaining the per patient exposure prediction matrix Ψ_*PID*_, the median TF alteration probability within each entity is compared against the alteration probability from the alteration detection pipeline.

## RESULTS AND DISCUSSION

### Impact of mutational signatures on transcription factor binding motifs

We computed the motif alteration signature using the conditional probability between the transcription factor alteration probability and the trinucleotide mutational signature as described in Eq. 5 and Eq. 6. The motif alteration results across all 512 JASPAR motifs are shown in Figure 3a for motif disruption (right) and motif creation (left). To capture similar patterns between transcription factor motifs, we applied a PAM clustering to the set of 512 motifs. The silhouette coefficient shows a local maximum at *k*_*clust*_ = 4 indicating that the optimal number of clusters is 4.

**Figure 3.**
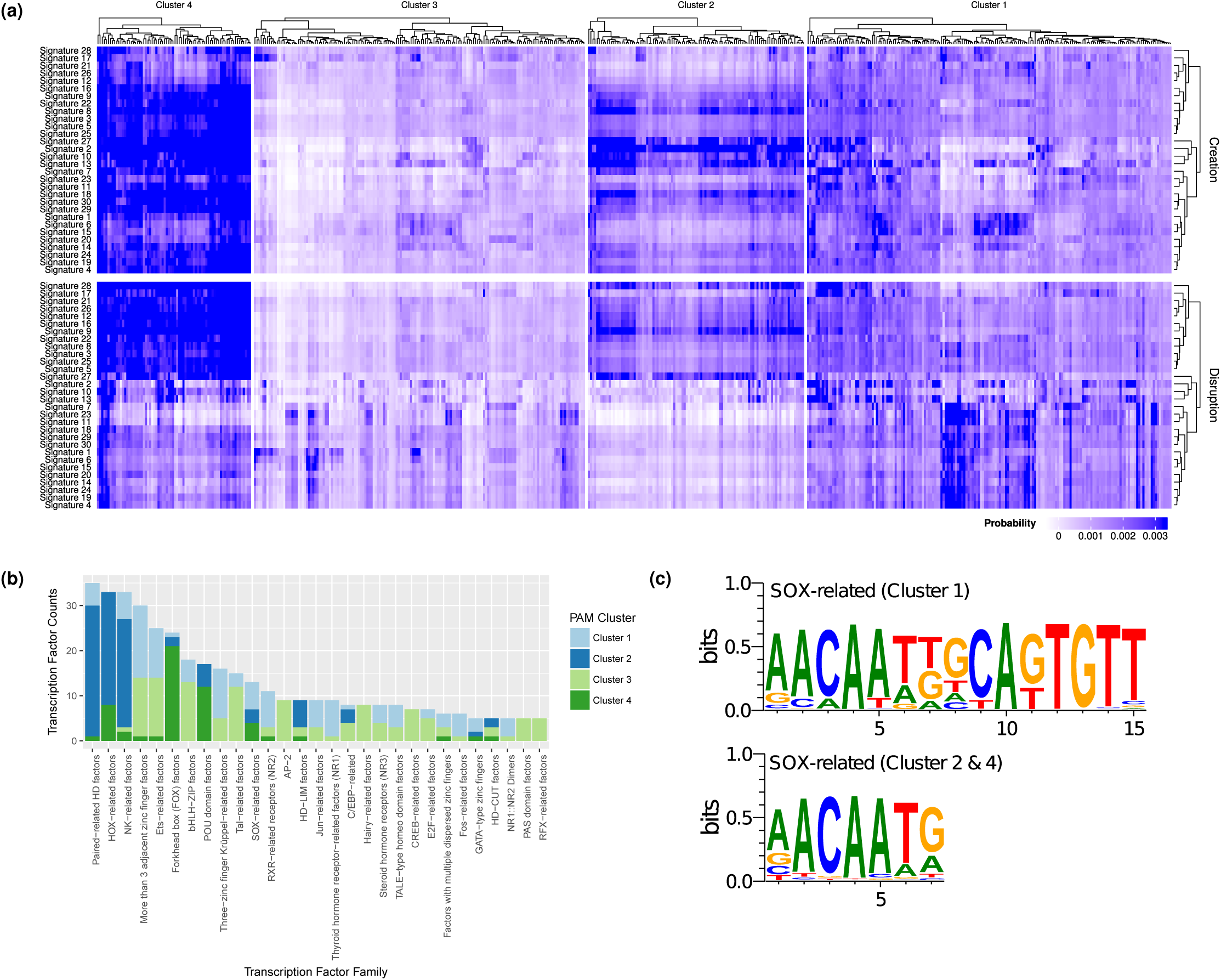
(a) Motif alteration signatures heatmap where the vertical axis corresponds to the 30 mutational signatures and the horizontal axis corresponds to the 512 motifs in JASPAR database. A *k*_*clust*_ = 4 PAM clustering performed over the TF alteration probability; (b) Cluster count statistics of each TF families with at least 5 members; (c) Familial binding profile of SOX-related motifs associated with PAM clusters.

The four clusters show a completely different behavior under the mutation signatures; whereas cluster 3 shows a low sensitivity to any of the mutational signatures (with some exceptions), cluster 4 on the other hand seems to be strongly impacted both by motif creation and disruption. Interestingly, comparing the disruption and creation heatmaps, we observe that the creation sensitivity for cluster 4 is high over all signatures, whereas we observe two groups of signatures for the disruption, with different impacts. We then studied the composition of TF motifs of each cluster. As expected, none of the clusters is dominated by a single TF family (Figure 3b). However, Figure 3b shows that some motif families are preferentially associated to one of the clusters. Forkhead motifs are in their vast majority associated with cluster 4 and show a high sensitivity to all mutational signatures. We also observe obvious mutual exclusivity between the cluster 1 and 2 versus cluster 3 and 4 for most of the TF families. SOX-related factors and C/EBP-related family, on the other hand, seem to be distributed across several clusters. For SOX-related motifs, we observe that these motifs form 2 major subgroups as shown in Figure 3c. SOX-related factors associated with PAM cluster 1 contain an extended binding sequence AACAATK**GCAKCAKTGTT** whereas those associated with PAM cluster 2 and 4 contain a shorter version AACAATG binding motif.

After this global analysis of the alteration patterns across all 512 motifs, we next focused on specific motifs related to transcription factors which play a role in cancer. We extracted from COSMIC cancer gene census the list of transcription factors to investigate how the binding motifs of transcription factors mutated in the cancer genome are affected by somatic mutations. In Figure 4, the motif alteration probability of 30 signatures are displayed with respect to the 94 TF motifs in this list. In this figure, the signatures are clustered according to the probability difference between creation and disruption. While most of the alterations appear to globally compensate each other (white shades in the third heatmap), some outliers arise in this view; for example, PRDM1 appears to have an excess of disruption in signatures 10, 13, 28 and 17. Interestingly, the PRDM1 gene itself is frequently mutated in diffuse large-cell B-cell lymphoma (DLBCL), a cancer type for which signature 17 is highly present. In particular, the four zinc-finger binding domains of PRDM1 show 5 positions at which there is a recurrent somatic mutation. Hence, in this cancer type, we have a coincidence of mutation in the DNA binding domain of the PRDM1 gene and high frequency of binding site alterations.

**Figure 4.**
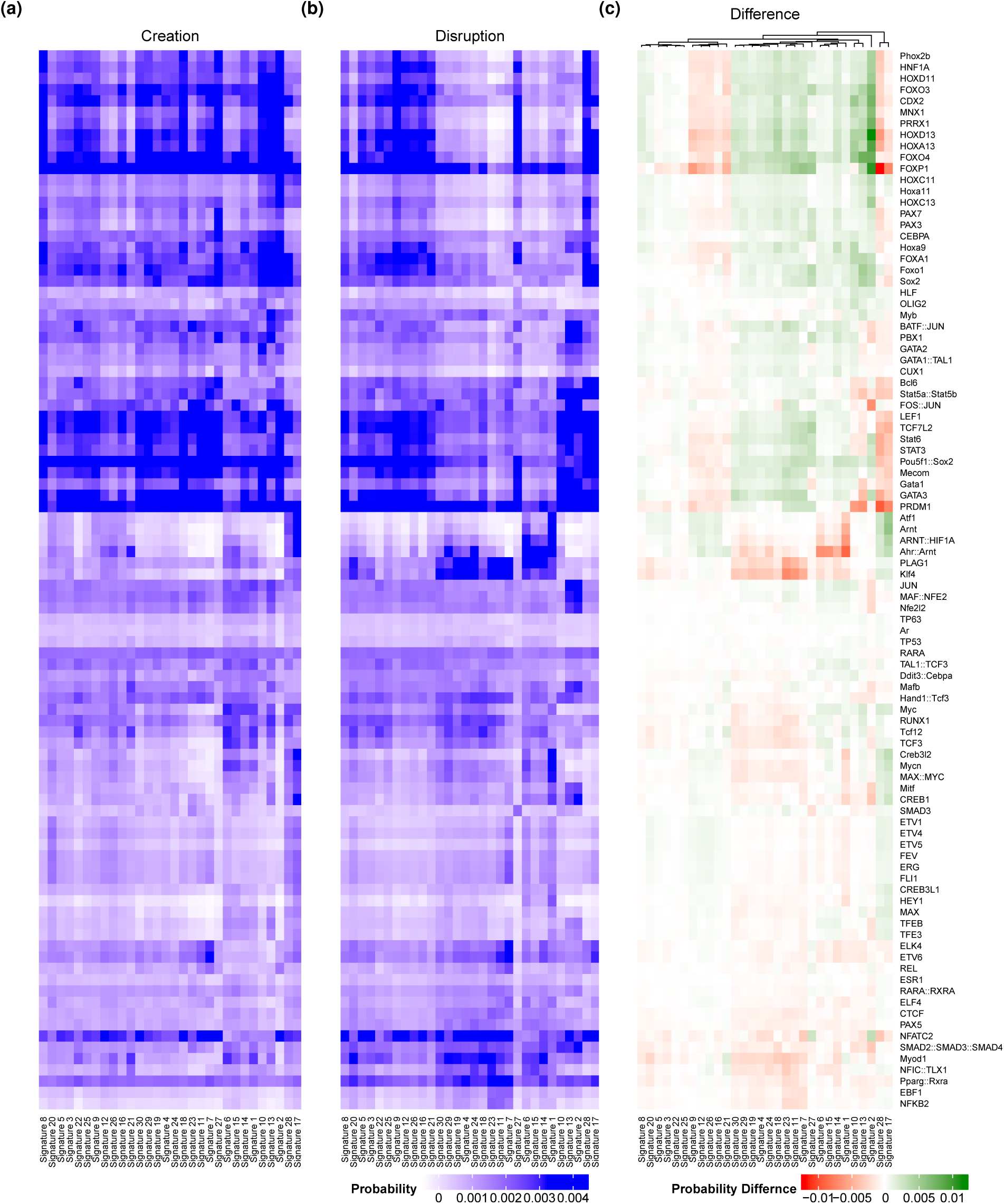
Motif alteration heatmap of TF in COSMIC cancer gene census (a) creation probabilities; (b) disruption probabilities; (c) probability difference (creation disruption).

In order to capture the complexity of the relation between the mutational signatures and the binding motifs, we applied SOM clustering to group motifs showing similar behaviors. We performed SOM clustering over the 192 possible mutational transitions (taking both creation and disruption into account). We also combined these mutational probabilities into the 30 mutational signatures.

The SOM clustering provides a clear picture on the similarity of the alteration behaviour of transcription factors across all trinucleotide mutational probability and the 30 mutational signatures. Transcription factors of the same transcription factor families often share similar binding sequence and their motif alteration behavior should also be similar. We observe that overall, transcription factors of the same family are well clustered together and globally share similar motif alteration probability pattern (Figure 5f and 5e). For example, the SOX transcription factor family is shown in Figure 5c and Figure 5d with SOX2, SOX4, SOX8, SOX9, SOX11, SOX10, SOX11, SOX17, and SOX21 clustered together. However, for specific signatures, differences between related transcription factors do exist. In Figure 5c, SOX10 appears to have a very similar creation probability to its neighbour SOX2 for signature 10, but shows a much lower creation probability in signature 23.

**Figure 5.**
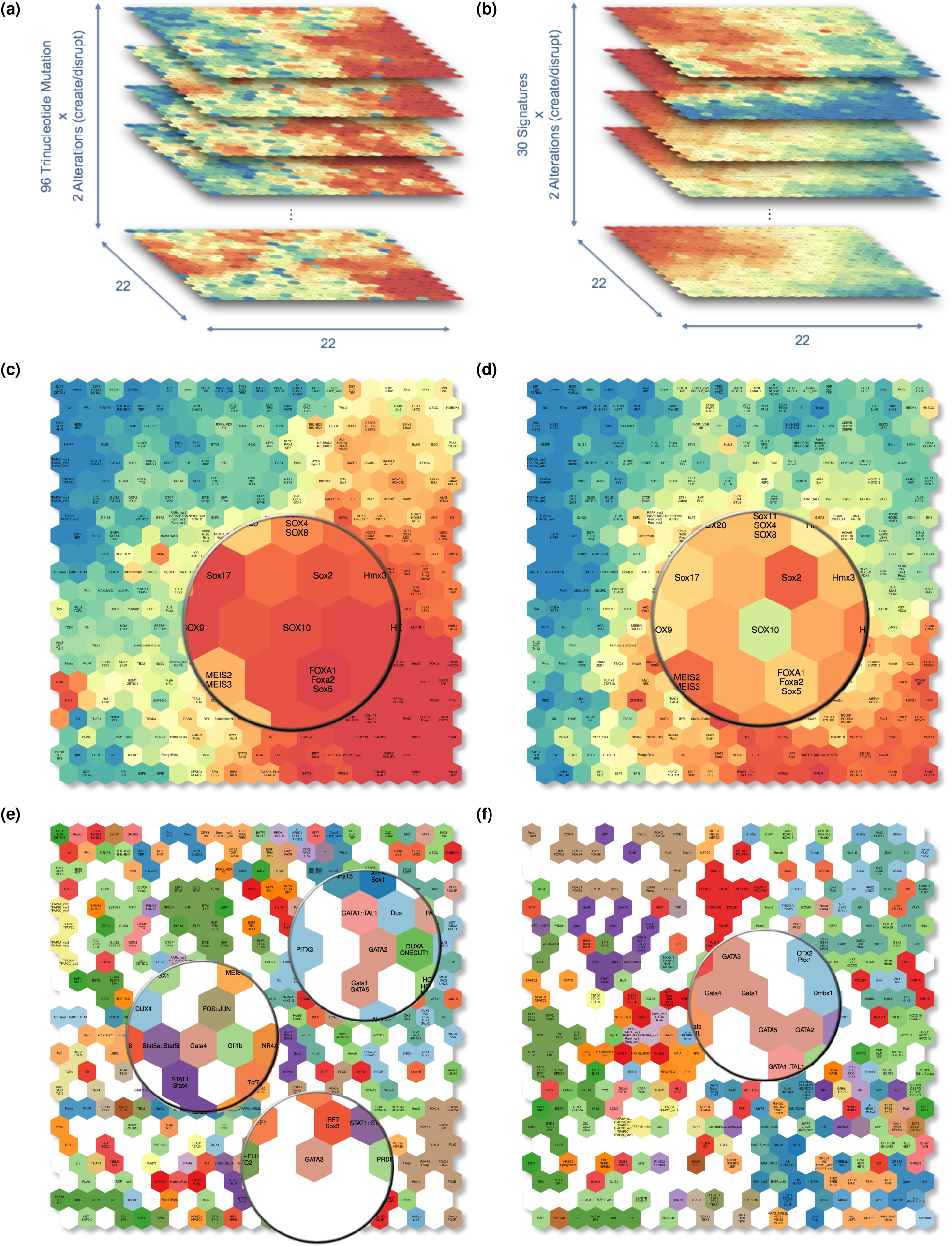
SOM clustering stack with (a) 192 variables of mutational probability (96 trinucleotide mutation types x 2 alteration types); (b) 60 variables of motif alteration signatures (30 mutational signatures x 2 alteration types); Transcription factor motifs of width 6 to 19: (c) SOM of Signature 10 creation probability; (d) SOM of Signature 23 motif creation probability; (e) SOM using of 30 motif alteration signatures with TF family colour coded; (f) SOM using 96 motif alteration probabilities with TF family colour coded.

Overall, by coloring the cells according to the family of the TF, we observe a global clustering of motifs from the same structural class (Figure 5e). However, this does not hold true for all families; we highlighted some members of the GATA-family in Figure 5e which appear to be dispersed across the SOM map. However, the SOM based on the 192 trinucleotide mutation probabilities (96 mutations for creation and disruption, Figure 5f) appears to group these motif together, highlighting the fact that mutational signatures are able to discriminate between related motifs.

### Correlation Between Motif Creation and Disruption Signature

The results from Figure 4 indicate that for many transcription factor, there is a compensating effect of creation and disruption. Further investigating this the relationship we found indeed that the alteration probability between creation and disruption event in all signatures have globally a strong correlation between the creation and disruption (Figure 6a). We found this correlation to be strongly related to the trinucleotide diversity of the motif matching sequences. To quantify this diversity, the trinucleotide content of each of the matching sequences is determined and the entropy is computed for each TF motif. The similarity between creation and disruption probability of each motif is evaluated using the absolute relative difference:

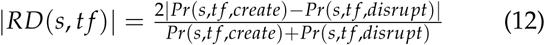

where *s* represents a signature and *t f* a binding motif. The scatter plot in Figure 6b shows an inverse relation between the complexity of the binding sequences (as measured by the entropy of these sequences) and the difference between creation and disruption. Hence, the more complex the motif content, the stronger the creation and the disruption signature probability are correlated.

**Figure 6.**
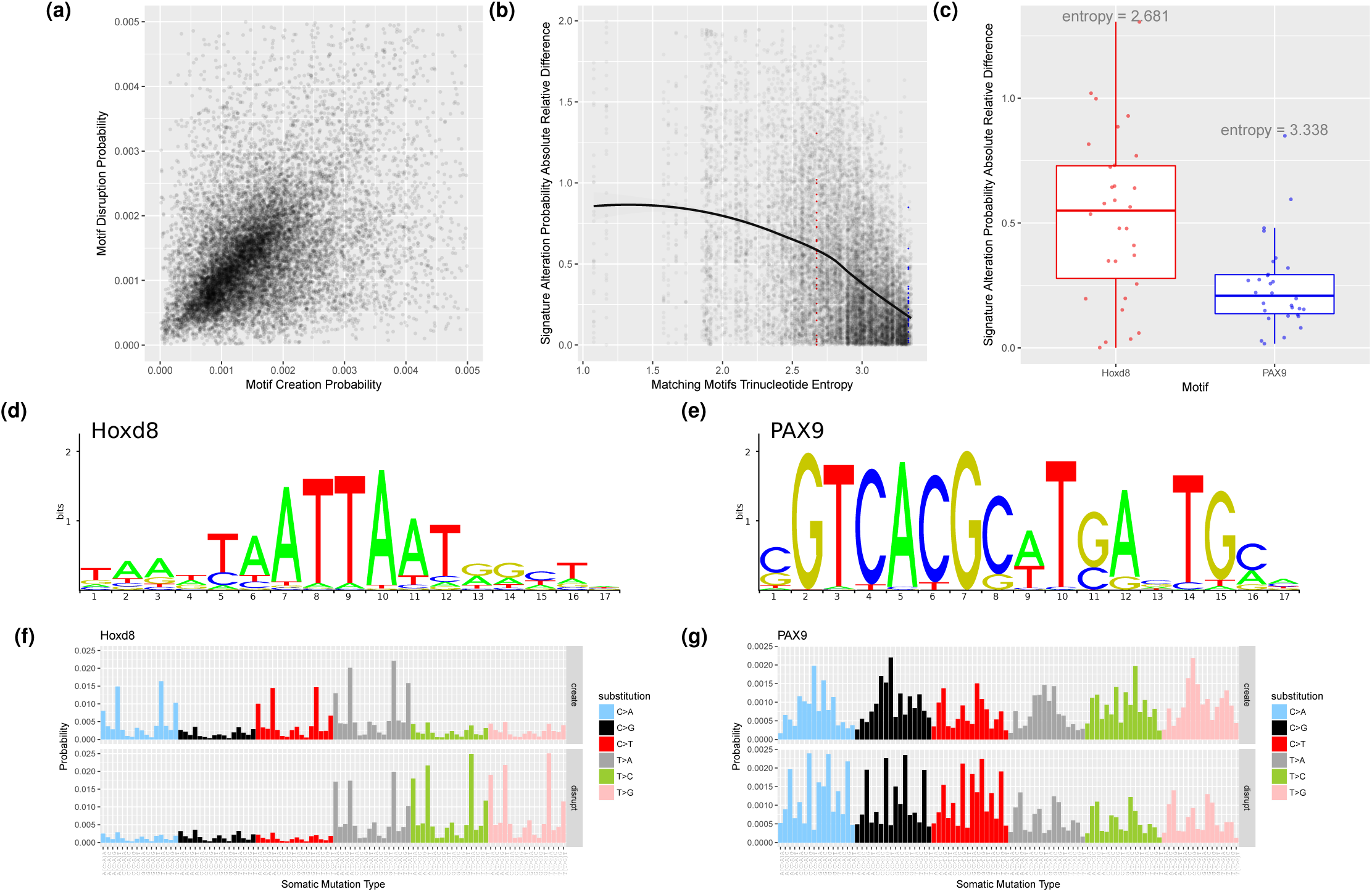
Demonstration of alteration correlation effect between creation and disruption probability: (a) Scatter plot of correlation between the creation and disruption probabilities for all 512 motifs across the 30 mutational signatures where each dot corresponds to a TF motif in 1 of the 30 signatures; Difference of creation/disruption probabilities vs. motif entropy, with Hoxd8 highlighted in red and Pax9 in blue; (c) Boxplot of alteration probability absolute relative difference of Hoxd8 versus PAX9; (d) PWM logo of Hoxd8; (e) PWM logo of PAX9; (f) Alteration probability of Hoxd8 with respect to 96 trinucleotides; (g) Alteration probability of PAX9 with respect to 96 trinucleotides.

This effect is illustrated using Hoxd8 and PAX9 as an example. To ensure that the motif width does not bias the results, both selected motifs have a width of 17 nucleotides. As illustrated in Figure 6d and Figure 6e, Hoxd8 and PAX9 have very different trinucleotide content. This is obvious looking at the motif logo. The motif alteration probability with respect to the 96 trinucleotide mutation is shown in Figure 6f. For Hoxd8, the motif alteration probability concentrates on trinucleotide mutation related to TAA, AAT, TTA, and ATT where the overlap of these trinucleotide mutation creation and disruption only occurs on A[T*>*A]A, A[T*>*A]T, T[T*>*A]A, and T[T*>*A]T.

On the opposite, the alteration probability of PAX9 is distributed along all the 96 trinucleotide mutations, resulting in the creation and disruption probability to be strongly correlated along the 96 trinucleotide mutations. This high similarity translates into a high similarity across the 30 mutational signatures. This is shown in Figure 6c for Hoxd8 and PAX9 across the 30 mutational signatures.

### Association of Deamination Signature and TFBS Creation

We next sought to validate our predictions using independent data. In [13], Zemojtel et al. described a set of transcription factors whose binding sites are frequently created as a result of CpG deamination process during evolution. These transcription factors include: C-Myc(Myc), Nfya, Nfyb, Oct4(POU5F1B), PAX5, Rxra, Usf1, and YY1. Given that some of the mutational signature in cancer are related to CpG deamination, we sought to verify if the same transcription factors are impacted by this process due to somatic mutations. The creation probability of these transcription factor motifs are plotted in Figure 7a over all 30 mutational signatures.

**Figure 7.**
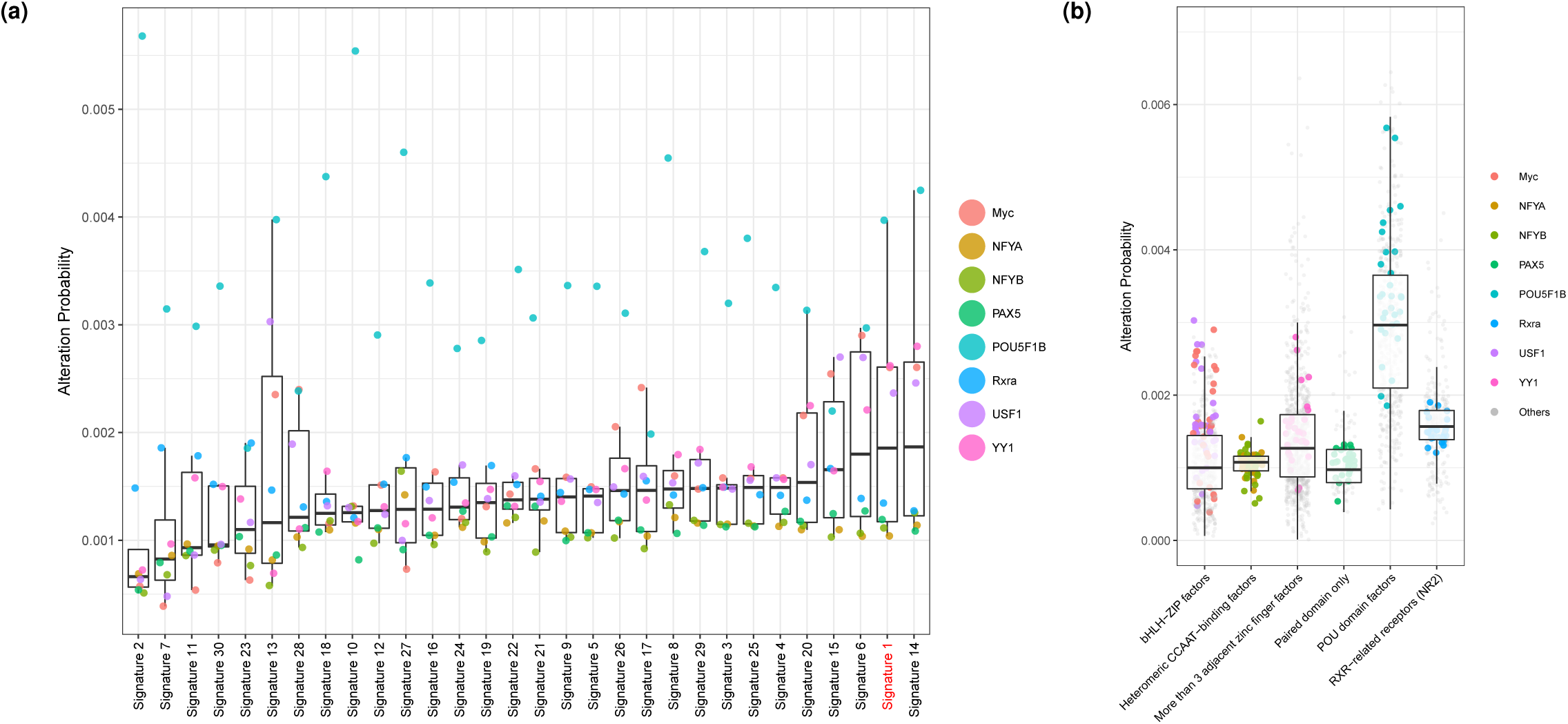
CpG deamination associated TF motif (a) creation probability for all 30 mutational signatures; (b) creation probability of all TF of the corresponding TF family.

Signature 1 was described as being related to deamination of 5-methylcytosine [14] and ranks as the second highest signature in terms of creation frequency of these motifs in Figure 7. Signature 14 shows a very similar mutational profile with strong C>T bias. In addition, it is interesting to note that the POU5F1B motif has a high creation probability compared to other TFs across all signatures. This phenomenon is due to the high genomic frequency of motifs which differ from POU5F1B binding motifs by one mutation, which gives rise to a large number of k-mers which closely resemble the POU5F1B binding domain and serve as a substrate for TF motif creation events.

If we extend this single TFs to the family they belong to, we also observe a much higher creation probability for the other members of the POU-family, compared to the families of the other impacted TFs (Figure 7b).

### Mutation associated Mechanisms

There are three main molecular mechanisms leading to single nucleotide mutations in cancer: i) defective DNA mismatch repair (MMR); ii) APOBEC activity; and iii) transcription-coupled nucleotide excision repair (NER). In the catalogue of mutational signatures, several signatures can be related to each of these processes. We wanted to investigate which transcription factor motifs are most impacted by these three different molecular mechanisms.

For each of these mechanisms, we summarized the alteration results for all signatures annotated to the same mechanism; APOBEC is related to signatures 2 and 13, MMR to signature 6,15,20 and 26, and NER to signatures 4,7,11 and 22. We displayed the differential alteration probabilities (creation disruption) in Figure 8a. Interestingly, a number of transcription factor motifs show reverse behaviors with respect to these three meachnisms. For example, Foxd3 shows a much higher creation probability for APOBEC and NER related signatures, whereas the opposite holds for MMR signatures.

**Figure 8.**
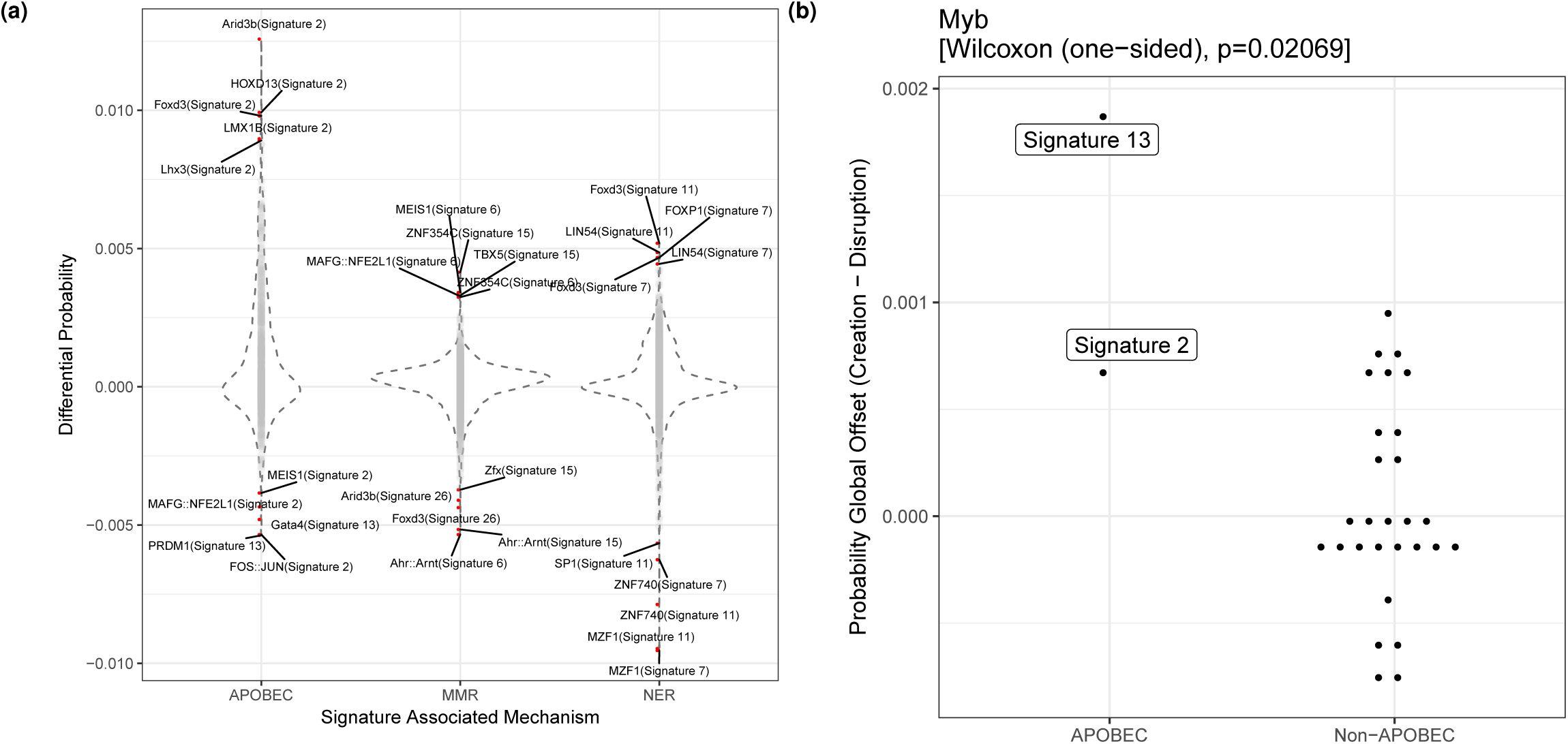
Multiple signatures driven molecular mechanism in cancer with associated TF and the corresponding global alteration offset comparison: (a) Top ranked TF and signature pair for each of the 3 associated mechanism; (b) APOBEC associated signatures and Myb.

As an example, it was shown that a mutation associated with the APOBEC signature leads to the creation of a Myb binding site [15]. To understand if this single event leading to a driver mutation might result from a more general impact of the APOBEC signature on the binding sites of Myb, we indeed observed that the two APOBEC signatures (signature 2 and 13) show the highest bias towards Myb motif creation compared to all other signatures (P-value = 0.02). Hence, this single driver event might result from an elevated creation rate of potential Myb binding site under these APOBEC signatures.

### Comparison with the TCGA Dataset

We then sought to validate our prediction using a dataset of observed SNV in cancer genomes. We used a previously published dataset of somatic mutations based on a set of 505 cancer genomes over 14 cancer types. The idea of the validation is to compare the predicted frequencies of motifs alteration that are explained purely by the mutational signature and its biases towards certain trinucleotides, with the frequencies of motif alteration observed when looking at a SNV dataset in cancer genomes. If the predicted and the observed frequencies are comparable, then most of the signal of motif creation or disruption can be explained by the impact of mutational signatures. If on the other hand we observe a difference, this discrepancy could be attributed to additional effects in the real dataset, such as positive or negative evolutionary pressure in the cancer dataset leading to an increased frequency of motif creation or disruption. Hence, our goal is to highlight such potential effects and to use our signature based prediction as a baseline.

In order to compute a predicted alteration probability per patient, we used the signature exposure of each patient, and performed a linear combination based on the exposure values. All patients related to a common cancer entity cohort (for example melanoma SKCM) are then combined. The differential alteration probability is computed for each motif and each cohort by first subtracting the motif disruption and creation probability for each TF and then taking the median of alteration difference by cancer entity and TF family. The result is shown in Figure 9.

**Figure 9.**
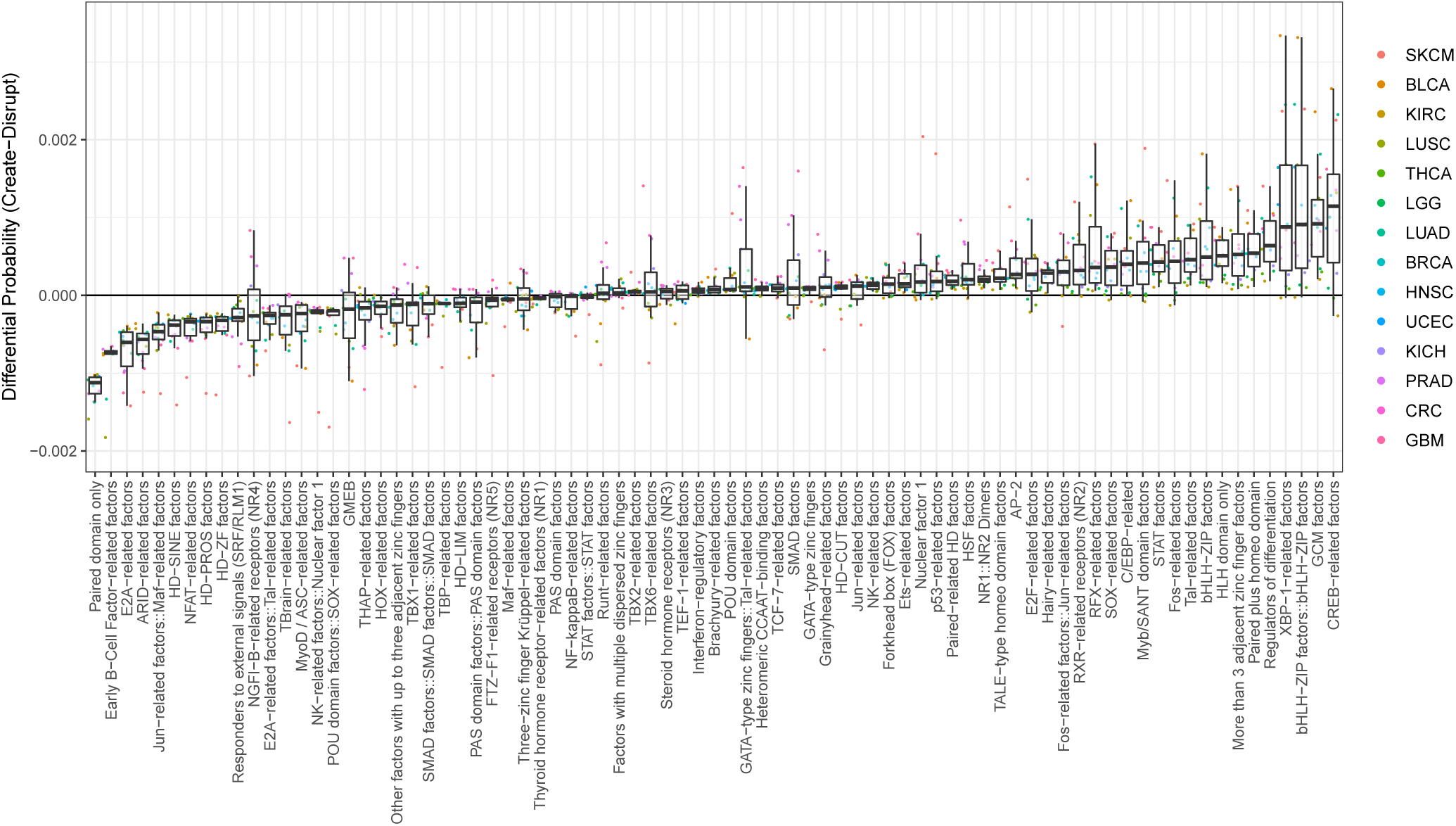
Inferred differential probability using exposure from the TCGA dataset. The differential probability is taken from the median of all TF members within the same family.

Interestingly, we observe three classes of TFs: TF families for which a positive alteration bias across all cancer types (for example motifs related to ETS factors or bZIP-related motifs) is observed, families for which this overall bias is generally negative (for example motifs related to NFAT factors) and those showing a mixed behavior. In this last category, we find SMAD related factors. Given the complexity of the TGF-*β* signalling processes in different cancer entities, the fact that their effector proteins of the SMAD family show different alteration probabilities across cancer types is plausible [16–18].

Next, we compared the prediction with the observed alterations across the TCGA dataset. To perform this comparison, we first determined the observed motif creation and disruption probabilities of all TFs across all samples within a given cohort using the motif alteration calling pipeline described in the method section. This pipeline predicts, for each SNV, whether it disrupts or creates potential binding motifs, and yields for each patient and each motif a creation and disruption probability. We then determined the Pearson correlation between these “observed” probabilities and the predicted probabilities from the mutational signature analysis over all samples of a cohort, combining creation and disruption. Hence, for each cohort, we obtain one correlation value. In all cases, the correlation value is positive and statistically significant. For some cancer entities, the correlation is low (*ρ <* 0.1 for lower grade glioma LGG), while it reaches *ρ* = 0.3 for the melanoma cohort. To explain the low correlation of some cancer entities, we compared this correlation value with the corresponding number of SNVs in the corresponding cancer entity in Figure 10. A linear regression is performed over the coefficient norm and the number of SNVs, and a significant correlation between the correlation norm and the number of SNVs is observed (*p* = 0.021). Given that our model is a theoretical prediction of the TFBS alterations probabilities across mutational signatures, we expect it to hold true in the limit of a growing number of SNVs. When the mutational load is smaller, additional effects such as selection or positional biases due to chromatin conformation contribute stronger to the pattern of mutations, explaining why the observed TFBS alteration patterns deviate from our theoretical model.

**Figure 10.**
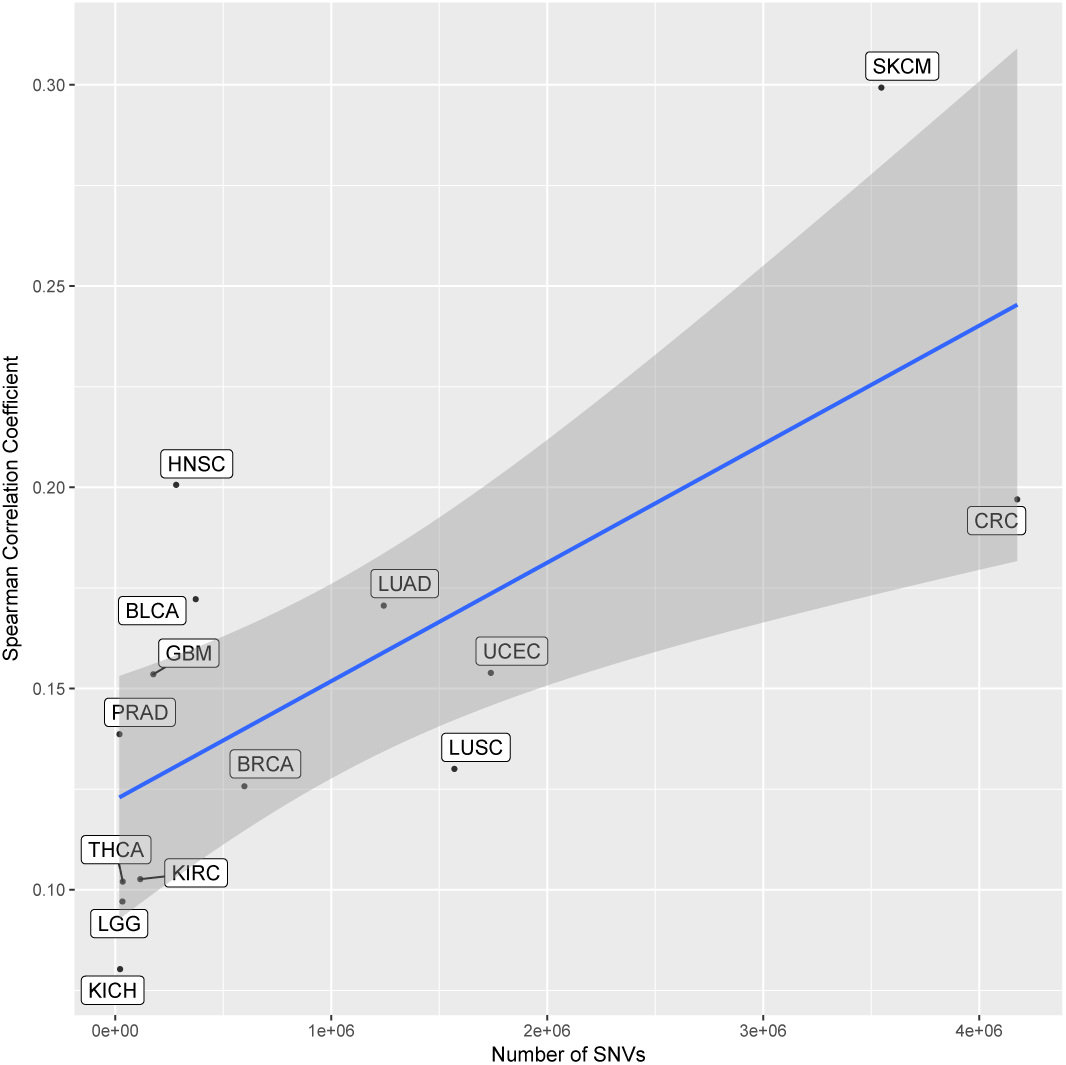
Comparison of cancer mutational signatures of transcription factor and TCGA dataset. Spearman correlation coefficient between the 1024 observed and signature inferred alteration probabilities (512 TFs x 2 alteration type) of each cancer entity. Linear regression between Spearman correlation and number of SNVs, p=2.1e-2.

## CONCLUSION

In this study, we investigated the theoretical impact of cancer mutational signatures on regulatory elements of the non-coding genome and provided an outline of a Bayesian framework for motif alteration analysis using mutational signatures. One of the key finding in this study is the correlation effect between motif creation and motif disruption. The correlation between the motif alteration probability was found to be strongly positively associated to the motif entropy. Further, previously described effects such as the impact of specific signatures on families of transcription factors can be reproduced by our theoretical model. An intriguing finding was that the described non-coding driver leading to Myb creation in T-ALL could be related to a global increased creation probability in APOBEC driven cancer types. Finally, the motif alteration signatures were used to infer the alteration events of each TCGA cohort using the corresponding signature exposure. It was found that the accuracy of prediction is positively correlated to the total number of SNVs within the cohort. Therefore, this motif alteration signature can serve as a background model for point mutation analysis for large dataset.

## ACKNOWLEDGEMENT

The authors would like to acknowledge the input and assistance from Jules Kerssemakers regarding the implementation of the data processing pipeline, and Ashwini Sharma for his scientific suggestions.

## APPENDIX

### TF PWM Alteration Probability Estimation Algorithm

To summarize, the alteration probability estimation algorithm consist of the following steps:

#### (1). Scan all k-mer sequences

~~~
For *i* = 1 to *I*, where *i* is the index of transcription factor set PWM
        Obtain *k*, where *k* is the width of pwm(*i*)
        For *p*_*k*_ = 1 to *P*_*k*_, where *p*_*k*_ is the index of k-mer sequence set SEQ
                Match pwm(*i*) against seq(*p*_*k*_)
                If p-value *>* 0.001
                        add seq(*p*_*k*_) to MatchSEQ set
~~~

#### (2). Counting every alteration event

~~~
For *i* = 1 to *I*, where *i* is the index of transcription factor set PWM
        For *j* = 1 to *J*, where *j* is the index of element in MatchSEQ
                For *q*_*trinuc*_ = 1 to *Q*_*trinuc*_, where *q*_*trinuc*_ is the index of the 96 mutation type
                        For *n*_*pos*_ = 1 to (*k -* 2), where *n*_*pos*_ is the position in sequence matchseq_*j*_
                        If matchseq [*n*_*pos*_ : *n*_*pos*_ + 2] == *mut*(*q*_*trinuc*_, *re f*)
                        mutseq = matchseq_*j*_
                        mutseq[*n*_*pos*_ : *n*_*pos*_ + 2] = *mut*(*q*_*trinuc*_, *alt*)
                        If mutseq not in matchseq_*j*_
~~~

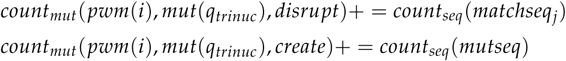

#### (3). Normalize probability

~~~
For *i* = 1 to *I*, where *i* is the index of transcription factor set PWM
        For *q*_*trinuc*_ = 1 to *Q*_*trinuc*_, where *Q*_*trinuc*_ is the index of the 96 mutation types
~~~

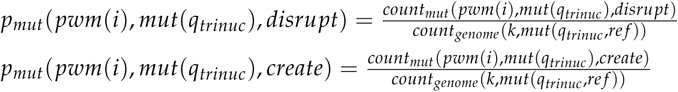

### Bayesian Inference of Transcription Factor Signature Alteration Probability

To compute the transcription factor signature alteration probability *Pr*(*a|s*_*i*_, *t f*_*k*_), we have:

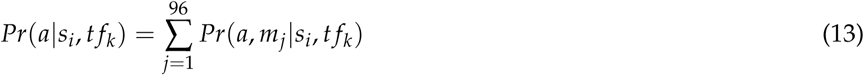

Based on the Bayesian tree described in Figure 1b, we have the joint probability of all parameters described by Eq. 14.

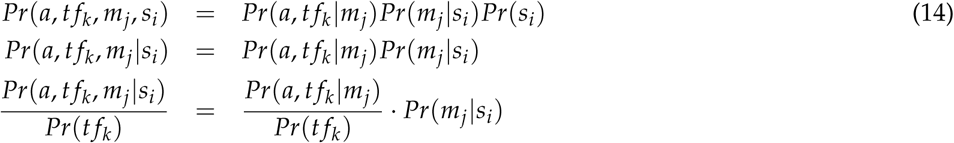

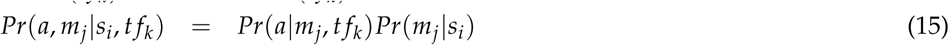

Combining Eq. 15 and Eq. 13, we have Eq. 5.

